# Tracking vaginal microbiome transitions in bacterial vaginosis for cues of antibiotic resilience

**DOI:** 10.64898/2026.04.28.721237

**Authors:** Apoorva Challa, S Afrin Zulfia, Anwesha Mohapatra, Sunil Nagpal, Mohammed Monzoorul Haque, Seema Sood, Garima Kachhawa, Geetha Thiagarajan, Bhupesh Taneja, Somesh Gupta

## Abstract

**Background:** Bacterial vaginosis (BV) is a common and difficult-to-treat vaginal disorder, with significant implications for reproductive health, particularly in low and middle-income countries. Clinical cure based on symptom resolution or Nugent scores often do not correspond to restoration of healthy vaginal microbiome. Factors underlying treatment failure remain poorly defined, warranting the need for understanding post-treatment microbiome dynamics to improve long-term outcomes.

**Objectives:** To delineate longitudinal vaginal bacteriome dynamics, integrating microbial composition, transition patterns, and clinical symptoms in a closely followed cohort of women with BV based on treatment outcome.

**Methods:** Vaginal swabs from reproductive-age women (18–45 years) were collected and classified as BV-positive (≥7) or healthy (≤3) using Nugent scoring. BV cases were treated using single-dose secnidazole and followed for three months. Sociodemographic, clinical, and behavioral data were statistically analyzed across groups. Vaginal microbiome composition was assessed using 16S rRNA sequencing, evaluating taxonomic profiles, alpha and beta diversity, differential abundance, and co-occurrence networks.

**Results:** Antibiotic treatment reduced overall microbial diversity and shifted community composition toward healthy controls, though relapse samples retained higher diversity of BV-associated taxa such as *Sneathia, Dialister*, and *Gardnerella*, while no-relapse and control groups showed higher *Lactobacillus* abundance. *Corynebacterium amycolatum* appeared protective, while *Mycoplasma* and *Fusobacterium* played symptom-specific roles. Microbial network analysis showed denser and more persistent associations in baseline and relapse groups, with *Sneathia* remaining a central node.

**Conclusion:** Short-term symptom resolution in BV does not correspond to full microbial recovery; necessary for long-term remission. Functional traits of resilient taxa like *Sneathia, Fannyhessea* and *Dialister* may confer resilience and enable recolonization, undermining long term treatment efficacy.

## Introduction

Bacterial vaginosis (BV) is the most prevalent vaginal disorder among women of reproductive age, characterized by a shift from a Lactobacillus-dominated ecosystem to a polymicrobial, anaerobe-rich community. Though often asymptomatic, BV is clinically significant due to its associations with increased susceptibility to sexually transmitted infections (STIs), pelvic inflammatory disease (PID), and adverse pregnancy outcomes such as preterm birth.^1-3^ Standard antibiotic regimens, including metronidazole, clindamycin, and secnidazole, achieve high short-term cure rates; however, recurrence affects up to 50-60% women within one year of treatment, often within the first three months.^4,5^ The prevalence is speculated to be much higher in the Global South, where, out of social stigma, a large fraction of cases remains unreported and untreated. ^6^

While Nugent based assessments currently guide the clinical diagnosis of BV, perturbations in total community dynamics of vaginal micro(b)environment can reveal a holistic picture of this disorder. Globally, the significance of lactobacilli dominated community state types (CSTs) in maintaining vaginal homeostasis have been prolifically reported.^7^ However, transitions in community composition and dynamics in response to antibiotic treatment remain poorly understood, especially from developing nations, like India. Handful of studies on BV recurrence report dominance of Mycoplasma, Veillonella, and Sneathia post-treatment along with re-emergence of Lactobacilli and persistence of recalcitrant polymicrobial biofilms - primarily structured by Gardnerella vaginalis.^8,9^ When carefully tracked for treatment outcomes, microbiome of relapse and recovery states may reveal important drivers of treatment-failure, especially when compared to their baseline (treatment naïve) state.

Hence, we sought to delineate longitudinal vaginal bacteriome dynamics, integrating microbial composition, transition patterns, and clinical symptoms in a closely followed cohort of women with BV based on treatment outcome.

## Materials and Methods

### Subject recruitment and sample collection

Vaginal swab samples were collected from the posterior fornix of women in the reproductive-age group of 18–45 years and were tested for BV using Nugent scoring method.^10^ Pregnant, lactating, menstruating women who had received antibiotics in the preceding four weeks or had a coexisting STI were excluded from the study. A Nugent score ≥ 7 was considered as BV-positive while those having a scores ≤ 3 and no characteristic symptoms of vaginitis (e.g. abnormal discharge, itching, burning, and lower abdominal pain) were considered controls (healthy controls). BV-positive women were treated with a single dose of oral secnidazole (2g), post which vaginal swab samples were recollected for Nugent scoring at day 7, day 30 and 3 months. Women negative for BV at day 7 but testing positive for BV either at day 30 or at 3 months were considered as relapse. Women who tested negative (Nugent score ≤ 3) for BV at day 7, day 30 and 3^rd^ month after treatment were categorized as no-relapse.

### Ethics Approval

The study was approved by the Institute Ethics Committee, All India Institute of Medical Sciences, New Delhi [IECPG-415/30.08.2018, RT-24/27.09.2018].

### Cohort biostatistics

Comparative analyses of metadata information with respect to socio-demographic, behavioural and clinical determinants were studied for four classes of samples namely a) BV-positive; (N=40) b) relapse (N=29) c) no-relapse (N=11) d) control (N=40). Descriptive statistical analysis was performed for all variables and frequency distributions were captured as values and percentages. Differential distribution analyses of continuous variables for the afore-mentioned groups were performed using Wilcoxon Rank-Sum (Mann-Whitney U) test. Correlation analysis between categorical concomitants was done using Fisher-exact test (Scipy, Python 3.10.16). Adjusted p-values were generated for the associated variables and a p-value of ≤ 0.05 was considered statistically significant.

### DNA extraction and 16S r RNA V3-V4 sequencing

Bacterial DNA was extracted and V3-V4 region of the 16S rRNA gene was amplified and sequencing was performed on a MiSeq V2 instrument (Illumina) as described earlier.^10^ A total of 0·2 million, 250 base pair (bp) paired end reads per sample were generated

### Microbiome analysis

#### a. Top Taxa

Identification of the top taxa across the 4 pre-defined classes based on the union of their sample-size-normalized taxonomic abundances was done using the phyloseq (version 1.50.0) and Microbiome (version 1.28.0) packages from R [version 4.4.2]. Top 10 taxa at OTU level along with top 5 taxa aggregated at genera and phyla levels were determined to observe taxonomic distribution among the classes.

#### b. Alpha Diversity

Diversity of microbes constituting a single sample was measured using alpha diversity indices such as Shannon (diversity), Inverse Simpson (evenness), Chao1 (richness) and Observed species (unique number of OTUs present in each group) and were computed using vegan package (version 2.6.10) from R. The indices were calculated on raw abundance data at OTU level. Comparison of alpha diversity indices across the four classes and statistical significance was calculated using Kruskal Wallis non-parametric test. Estimation of significant change in alpha diversity between baseline and post-treatment samples was computed using paired analysis signed rank test. Post-hoc tests were done using the Bonferroni correction method. For visualization, violin plots and paired samples between baseline and post-treatment (relapse and no-relapse) were generated using the ggviolin() and ggpaired() functions respectively from the ggpubr R package.

#### c. Beta Diversity

Variation in microbial species composition of samples belonging to the different classes (i.e. beta diversity measure) was assessed on relative taxonomic abundance data. Bray-Curtis distance between datapoints was computed by vegdist() function from vegan package and the clusters were further visualized in a reduced dimensional space using ordinate() function from phyloseq package. Plots for visualization of the clusters were generated using ggplot2 and ggpubr. Homogeneity in the spread of samples within classes was analyzed using betadisper() from vegan package in R. To test for compositional shift in the classes due to various experimental parameters (e.g. age, previous treatment, group), permutational multivariate ANOVA (PERMANOVA) was performed using adonis2(), where variance per predictor was sequentially tested by=“terms”, from pairwise Adonis (version 0.4.1) package of R. Visualization plots, depicting movement of erstwhile baseline samples to post treatment relapse or no-relapse groups were created using generate_beta_ordination_pair() function from MicrobiomeStat package (version 1.2).

#### d. Differential Abundance Analysis

Differential taxa between groups based on microbial count data was analysed using ancombc2() function from ANCOMBC (version 2.8.1) package in R, which analyses microbial composition by accounting and correcting for sampling fraction and taxon-specific biases. The function also generated a sensitivity score to overcome bias introduced due to pseudo-count addition to log-transformed raw abundance count data. Parameters used for the function included prevalence cut-off of 0·1 (10%), alpha significance threshold of 0·05 and adjustment of p-value via Benjamini Hochberg method. Furthermore, covariates (e.g. age, previous treatment) having potential influence on taxonomic abundance were also included along-with ‘group’ factor in metadata. The taxa passing both significance (adjusted p-value) and sensitivity thresholds were considered differentially abundant and their corresponding effect sizes (log fold change values) were visualized as lefse bar plots, generated using ggplot2 package in R.

#### e. Co-occurrence Analysis

The microbial co-occurrence network for each group was constructed using MetagenoNets tool.^11^ The input abundance data was transformed using the centered log-ratio (CLR) method. The default algorithm and prevalence cut-off were applied, with an occurrence cut-off set at 0·2 and p-value significance level of 0·05. To generate the Venn diagram of shared edges and nodes between groups and create shared interaction network, the edge list derived from the previously generated co-occurrence networks were given as input to Netsets.js and GraphOnline tool, respectively.^12,13^ Cytoscape was used to study network dynamics and fluctuations of individual BV associated organisms across the analysed cohorts.^14^

## Results

### Cohort demographic and clinical statistics

Average age of women who relapsed was marginally higher (31·1 ± 7·07) than those who did not relapse (28·8 ± 7·76). However, age, marital status, sexual activity and history of infertility were not found to be associated with treatment outcome. The demographic characteristics of BV positive women against healthy controls were described (Table 1). ^10^

While the number of women who did not relapse (no-relapse) was small, underpowering the statistical inferences, a noticeable reduction in symptoms such as itching, burning, dysuria, dyspareunia, lower abdominal pain, odour and burning micturition was apparent among them. (Table 2). On an aggregate symptom scale, one or more symptoms appeared to persist among women treated for BV suggesting that antibiotic treatment did not completely alleviate their symptomatic profile (Table 2). Even though the recovery group experienced a drop in Nugent scores (i.e. ≥ 7 to ≤ 3), 70% women still exhibited Amsel criteria consistent with BV. Next, we probed the impact of antibiotic treatment on the vaginal bacteriome.

### Antibiotic treatment attempts re-stabilisation of alpha diversity and the dominant vaginal flora

In treatment-naïve women, antibiotic therapy significantly reduced vaginal bacteriome alpha diversity, regardless of treatment outcome (Figure 1a-h; Figure S1a-h) (Kruskal-Wallis test, Bonferroni p-value < 0·01). However, diversity metrices of women experiencing relapse were significantly higher than both no-relapse and control groups (Figure 1b-d, f-h; Figure S1b-d, f-h). Considering control group as a healthy reference, an incremental recovery of alpha diversity post-antibiotic treatment was apparent. This suggested the putative resilience of certain bacteria in the relapse group that resisted re-stabilisation towards a healthy community.

**Fig. 1.**
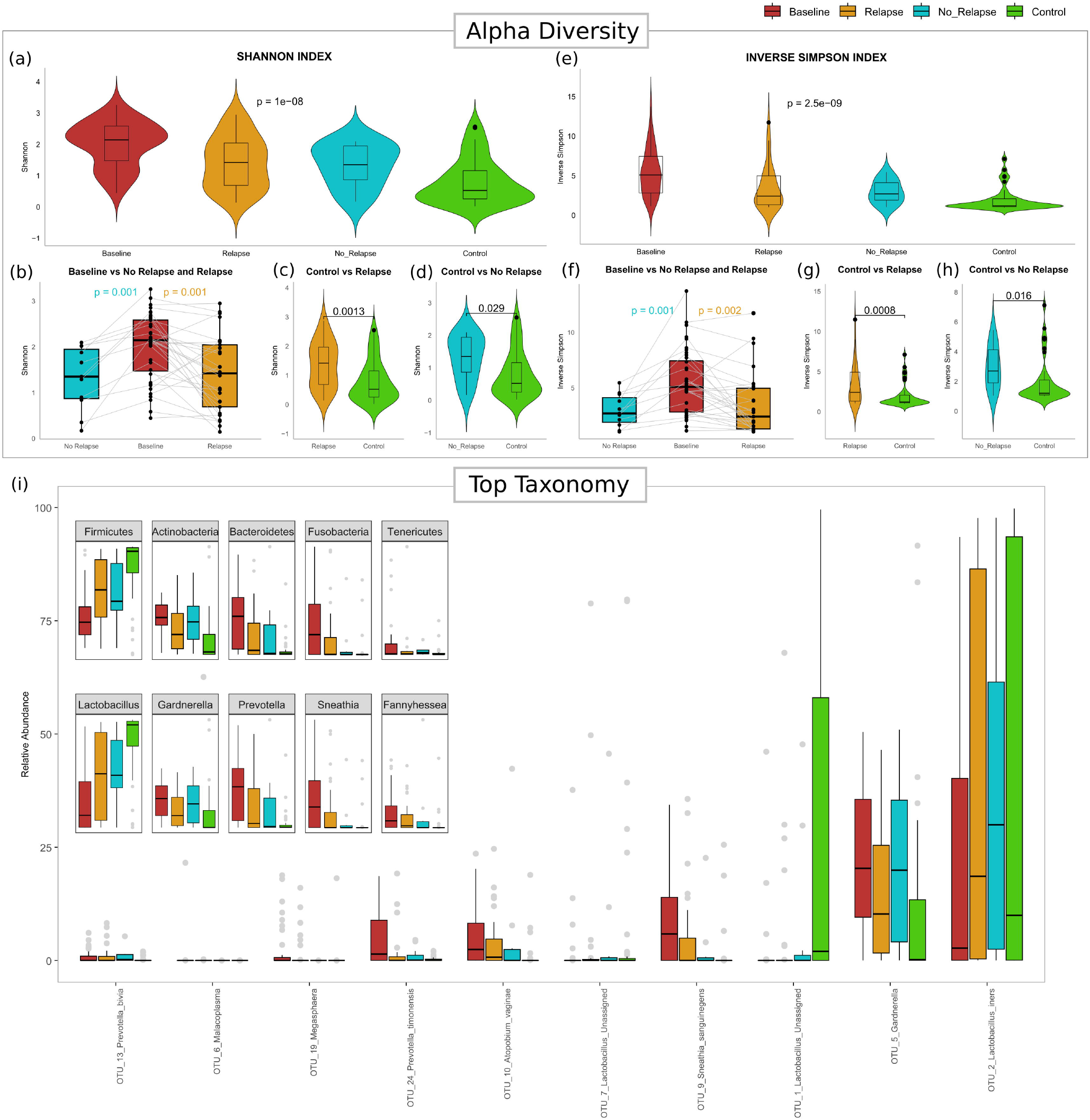

Tracing the composition of predominant flora, an incremental increase in abundance of phylum Firmicutes and decline in phyla Bacteroidetes, Fusobacteria and Tenericutes could be observed across the four groups (inset Figure 1i). Bacteroidetes and Fusobacteria were further observed to have higher abundance in relapse group than no-relapse group, indicating the putative phylum affiliation of resilient bacteria that may drive recurrence. At the genus level, an increase in the abundance of *Lactobacillus* (phylum: Firmicutes) and decline in abundance of *Prevotella* (phylum: Bacteroidetes), *Sneathia* (phylum: Fusobacteria) and *Fannyhessea* (previously *Atopobium*, phylum: Actinobacteria), potentially linked to resilience was evident by their higher abundance in relapse than no-relapse group. Despite, the overall depletion of OTU_10 (*Fannyhessea vaginae*) and OTU_9 (*Sneathia sanguinegens*), these species were also found to be higher in abundance among women who relapsed after treatment than those who did not. Interestingly, while phyla Actinobacteria and Tenericutes, genus *Gardnerella*, OTU_13 (*Prevotella bivia*), OTU_24 (*Prevotella timonensis*) and OTU_5 (*Gardnerella*) also displayed an overall decline post treatment in comparison to baseline samples, the incremental changes in their abundance were inconsistent.

With cues of re-stabilisation and resilience of alpha diversity and dominant flora post-treatment, we next probed its impact on inter-sample diversity and total community structure.

### Dissimilarities in bacteriome composition reveal signs of recovery and resilience

Bray-Curtis dissimilarities among BV-positive women were smaller, as indicated by close spatial clustering, compared to controls (Figure 2a, b; Figure S1i, j). Treatment increased inter-sample dissimilarity, shifting samples closer to control state. While relapse samples clustered close to baseline with limited increase in inter-sample dissimilarity, the no-relapse group with high dissimilarity, aligned closer to controls. These observations reinforce resistance to antibiotic treatment among women who relapse. Age and previous treatment were accounted for while analysing beta-diversity differences.

**Fig. 2.**
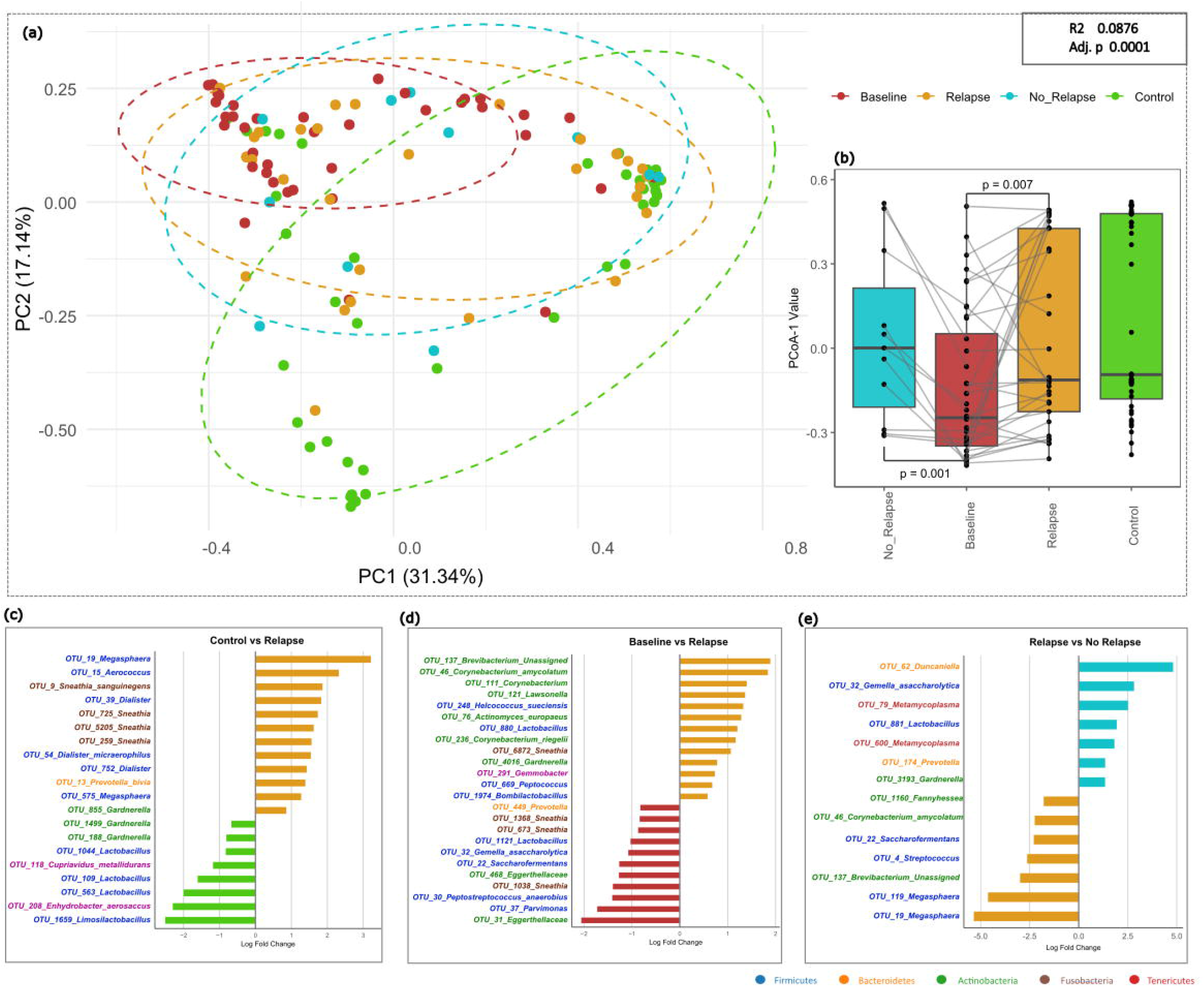

To identify key taxa driving treatment outcomes, differential abundance (DA) analysis was performed by correcting for age and previous treatment. *Lactobacillus* was significantly higher in no-relapse and control groups, as compared to women who relapsed (Figure 2c, d, e). Contrarily, enrichment of *Dialister* and *Sneathia* was observed in baseline and relapse groups when compared to women who did not relapse (Figure 2c, d, e). Notably, on comparing relapse and control groups, more than two-fold higher abundance of *Megasphaera, Aerococcus, Sneathia sanguinegens* and *Dialister*, along with multiple OTUs of *Sneathia* was observed in the former (Figure 2c). Interestingly, *Megasphaera* was enriched multi-fold in relapse than no-relapse group. Details regarding effect-size/log fold change, taxon involved, significance values etc., are described in Table ST1.

These observations raised the next pertinent question – how does antibiotic treatment shape community network dynamics i.e. microbe-microbe co-occurrence relationships?

### Vaginal microenvironment may carry symptom-specific bacteriome signatures

Given the broad but overlapping symptom profile in BV, we analysed symptom-microbe associations, examining non-collinear pairs of symptoms individually and omitting collinear counterparts for linear modelling of ANCOMBC2 to adjust for confounders. Marked enrichment of *Mobiluncus mulieris*, key for Nugent scoring, was observed in patients reporting abnormal discharge and dysuria. Another proinflammatory bacterium *Metamycoplasma* was also elevated in patients with abnormal discharge and several other symptoms including itching, dyspareunia, and malodour. Genital burning was associated with multi-fold increase in *Mycoplasma hominis*, while its absence was linked to significantly higher abundance of *Veillonella montpellierensis*. Malodour was associated with elevated levels of *Fusobacterium*.

### Tracking shifts in community networks

As reported previously, the BV community network was densely connected (234 edges) as compared to co-occurrence patterns post-treatment (relapse: 156 edges, no relapse: 54 edges) and control (116 edges) groups.^10^ This transition of network densities towards a sparse network after antibiotic treatment provides initial evidence of community restructuring towards a healthy state.

At the community level, despite correcting for network edge density, *Sneathia sanguinegens* persisted as a key regulatory node with highest degree transitions from baseline to relapse networks (Figure 3a-d). Its negative association with *Lactobacillus* species was additionally apparent in both states. Furthermore, multiple nodes pertaining to *Gardnerella* were prevalent among top degree nodes in both baseline and relapse networks. Noteworthy was the emergence of increased and positive associations of *Corynebacterium amycolatum* post-treatment, while *Sneathia* lost majority of its edges in no-relapse as compared to both baseline and relapse networks.

**Fig. 3.**
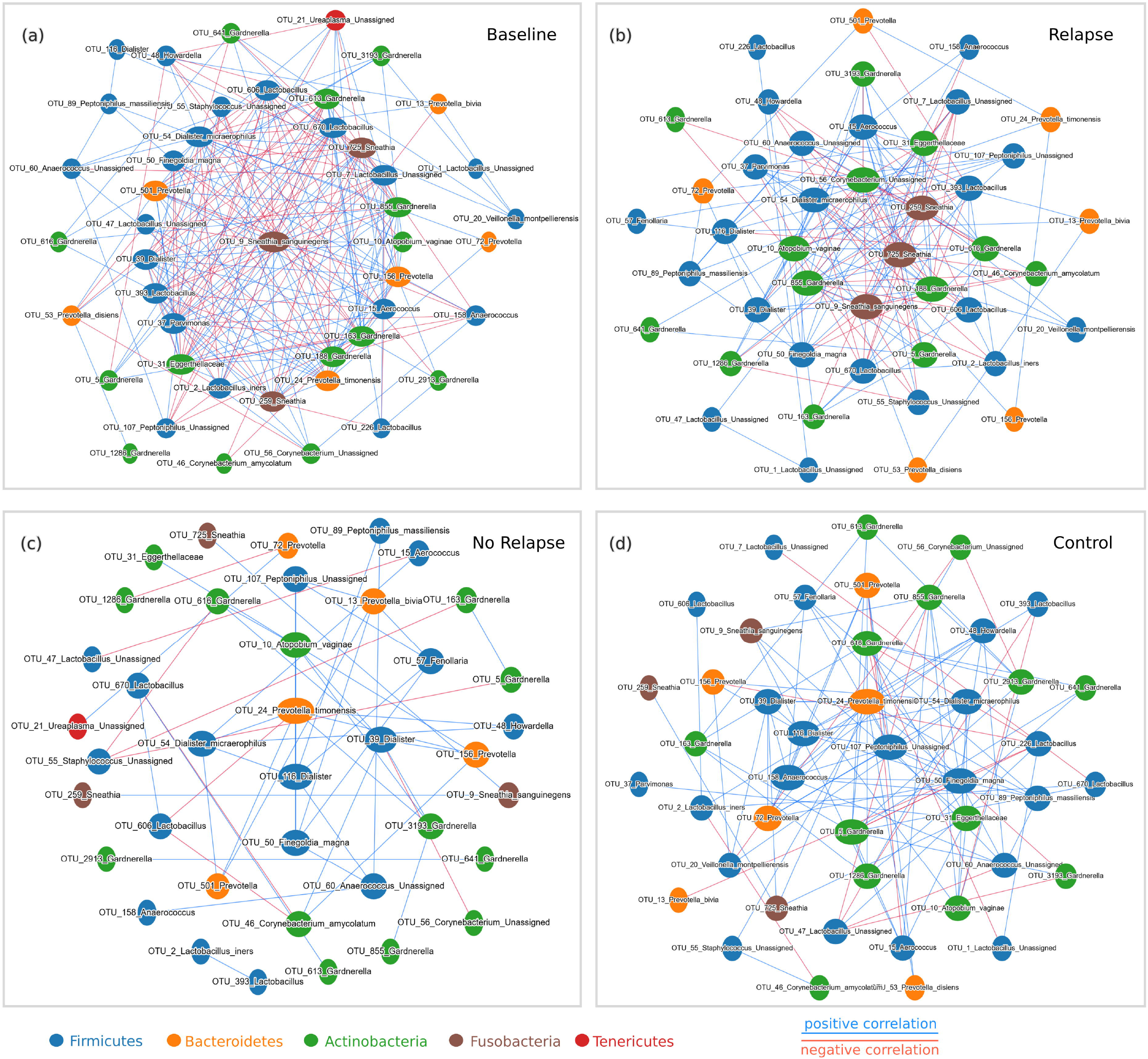

Comparison of the node and edge compositions further revealed that while baseline and relapse group exclusively shared 48 associations, these reduced to just 5 common edges between baseline and corresponding no-relapse groups. (Figure 4a-e). The overlap in edge compositions of baseline and relapse groups was primarily contributed by sustained associations of *Sneathia, Gardnerella* and *Dialister* OTUs (Figure 4d). Notably, five associations, pertaining to *Prevotella timonensis* with *Dialister microaerophilus, Anaerococcus* spp. with *Peptoniphilus massillensis*, and mutual associations of *Lactobacilli* and *Gardnerella* OTUs, were observed as core interactions across all four groups (Figure 4c).

**Fig. 4.**
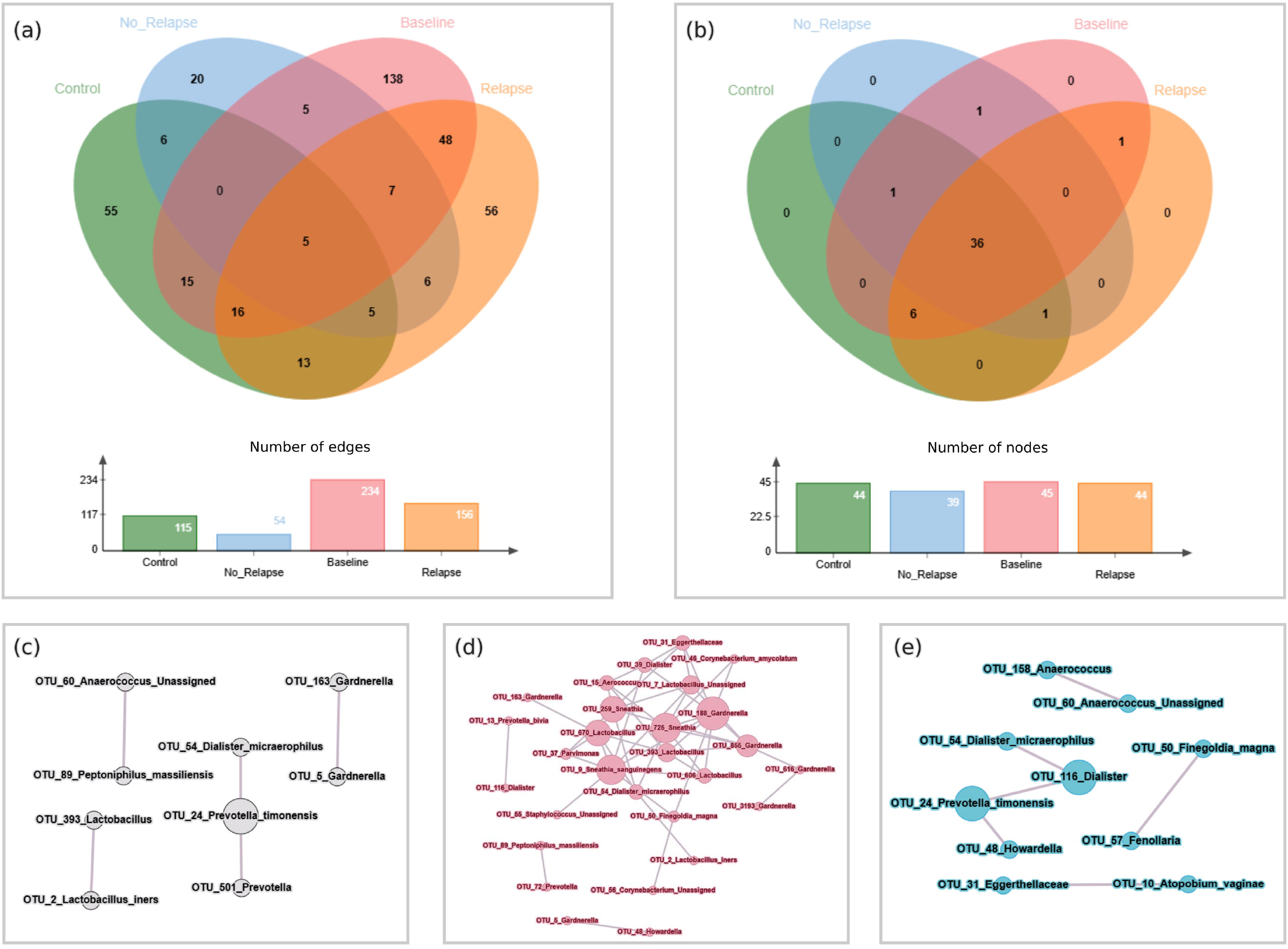

Examination of co-occurring relationships of *Sneathia* (Figure 5a-d) between baseline vs relapse, and baseline vs no-relapse reinforced the aforementioned interpretations. Sustenance of positive associations of *Sneathia* with *Gardnerella, Dialister, Aerococcus*; and negative associations with *Lactobacilli* and *C. amycolatum* between baseline and relapse samples were observed but markedly disrupted in no-relapse group. While *Sneathia’s* positive (and even negative) associations were disrupted, the mutually positive associations among *Sneathia* OTUs persisted. This indicates a potential gap in eradicating *Sneathia*, and may explain the >80% relapse rate for BV during the 1-year follow-up period. (*10*) Additionally, we present transitions specific to *Lactobacillus* (Figure S3), *Gardnerella* (Figure S4) and *Corynebacterium* (Figure S5) across the studied groups to further aid the interpretation of these community dynamics.

**Fig. 5.**
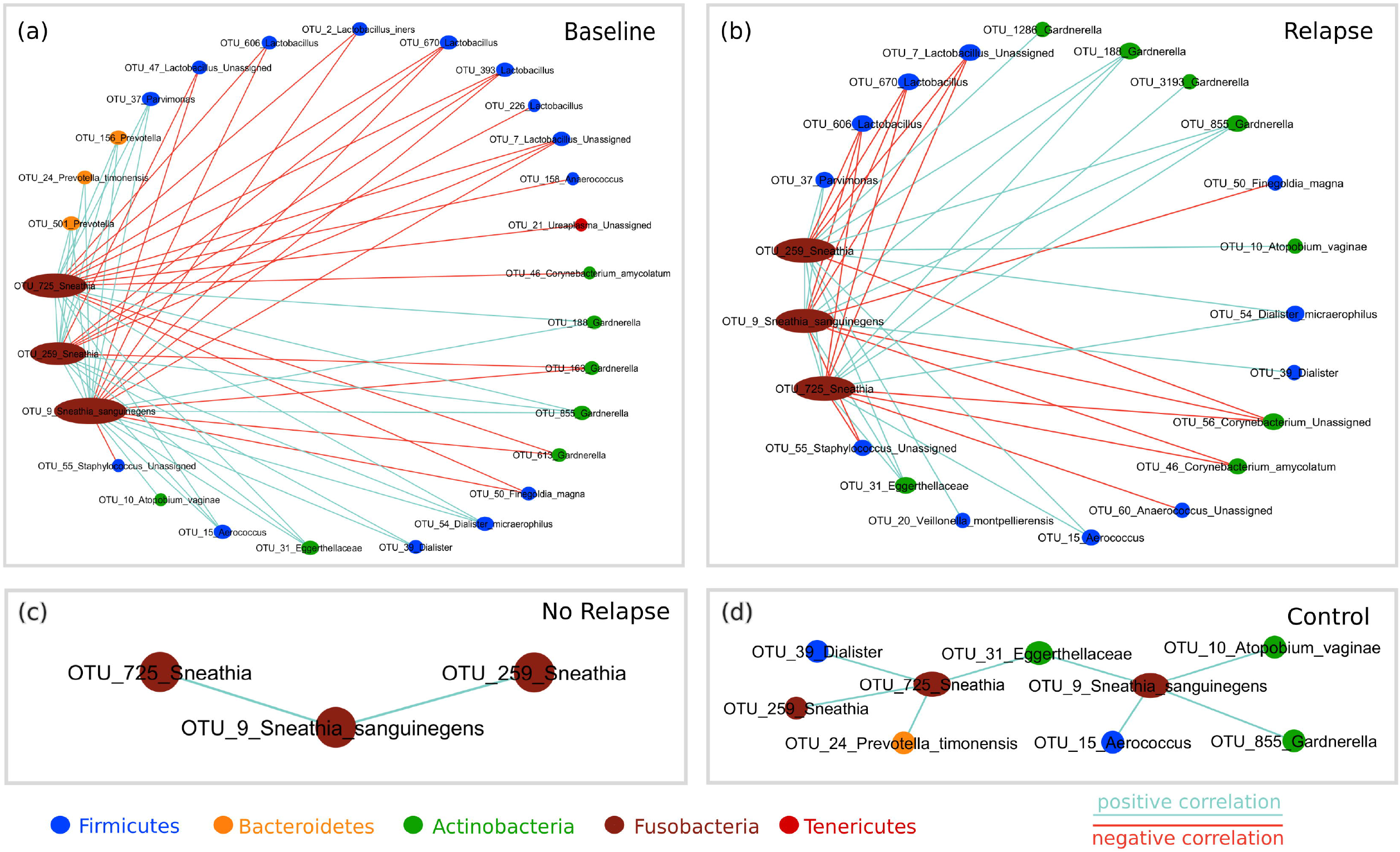

Collectively, these dynamic transitions highlight the consistent potential role of *Sneathia* as a key regulatory member of vaginal bacteriome in BV wherein disruptions may facilitate potential for recovery of community network towards a healthier state. Additionally, while corroborating the well-founded representation of *Gardnerella* and *Lactobacilli*, these dynamics also revealed previously underreported presence of *Corynebacterium amycolatum* as an additional player in vaginal community, warranting further investigation of its prognostic and therapeutic role in BV.

## Discussion

Despite the high rate of recurrence and complex vaginal microbial interplay, BV diagnosis and treatment landscape have not evolved beyond microscopic/symptomatic scores and broad-spectrum antibiotic administration. While, cross-sectional studies have advanced our understanding of BV, longitudinal studies-particularly from the Global South, remain scarce. Despite a modest sample size, our study addresses this gap by providing longitudinal insights into the vaginal microbiome structure and dynamics in secnidazole treated India women with BV.

In comparison to metronidazole (half-life 8 hours) widely reported in previous studies, secnidazole exhibits higher activity and prolonged bioavailability (half-life 17 hours) with similar relapse rates. ^15^ At 1-month follow-up, more than 50% women relapsed and by the end of 6-months, relapse rates increased to > 80%, consistent with previous findings (Table 1). Although 58% of women who relapsed were symptomatic, BV recurrence in asymptomatic women underscores the need to analyse the underlying microbiome structure that influences recovery. Clinical evidence suggests that BV frequently relapses with sufficient follow-up, emphasizing the need to characterize microbiome dynamics relative to both treatment naïve and healthy individuals.

### Impact of antibiotic treatment on the recovery of microbial diversity and the core microflora of Sneathia, Dialister, Fannyhessea, Gardnerella and Lactobacillus

While antibiotic therapy did alleviate overt symptoms, complete restoration of microbial homeostasis relative to controls was rarely achieved (Figure 2). Treatment reduced microbial diversity towards control state (Figure 2); however, it was still higher in both relapse and no-relapse groups. Interestingly, the no-relapse group exhibited lower diversity than relapse group, indicating a partial recovery of native microflora (Figure 2). Previous studies concur that recurrent BV is marked by a high alpha diversity and a failure to restore a low-diversity Lactobacillus-flora.^16-18^

Microbial composition analysis revealed a steady increase in abundance of Firmicutes phylum (genus *Lactobacillus*) from treatment-naïve to healthy state and, no relapse cases, suggesting shift towards healthy environment(Figure 2).^19,20^ Gradual decline in Fusobacteria, Bacteroidetes and Actinobacteria (Actinomycetota), typically enriched in BV vs post-treatment, with higher abundance in relapse vs no-relapse cases suggests their transitionary state but potential role in persistence of infection. Similarly, lower levels of Lactobacillus (phylum Firmicutes) in no-relapse and, persistence of Sneathia, the opportunistic pathobiont Dialister, and Fannyhessea (phylum Actinobacteria) was notable for relapse group (Figure 2) Presence of *Gardnerella* - a hallmark organism of BV, was noted in all groups, however with lower abundance in healthy samples.^21^

While persistence of *Sneathia* spp. has previously been reported to promote chronic inflammation and impairment of epithelial integrity, *Fannyhessea* spp. is overrepresented in BV recurrence owing to its association with biofilms.^22,23^ *Dialister*, on the other hand have been observed resistant to various antibiotics including metronidazole, macrolides, and fluoroquinolones.^24^ Clustering of samples based on microbial composition, indicated that relapse samples clustered closer to treatment-naïve cohort, while no-relapse samples were closer to healthy controls (Figure 3).While *Lactobacillus* was differentially abundant in healthy and no-relapse groups, *Dialister* and *Sneathia* were enriched in BV and relapse groups. Compared to no-relapse group, *Megasphaera* was multi-fold enriched in women who relapsed. *Megasphaera* has previously been associated with recurrent BV, especially if abundant in the pre-antibiotic treated samples. ^25,26^

### Symptom-microbe association: contribution of Mobiluncus, Mycoplasma and Fusobacterium in BV

Symptom-specific microbial associations added valuable insights into the role of *Mobiluncus mulieris, Mycoplasma hominis*, and *Metamycoplasma* in BV symptomology (Table ST2). These rather underdiscussed bacteria are known to elevate inflammatory responses and epithelial disruption, implicating their presence to an aggravated state, reflected in apparent symptoms (Table 2). *M. hominis*, found enriched in women reporting with burning sensation is rather known to colonize vaginal mucosa, breach the epithelial barrier, triggering cellular and DNA damage pathways.^27,28^ Members of *Fusobacterium*, observed elevated in women reporting vaginal malodour in this study, are similarly known to produce ammonia contributing to high vaginal pH, characteristic malodour, and biofilm maturation.^29^

### Community network analysis revealed microbial community transitions linked with treatment outcome

The incremental shifts in community networks guided intuitive and interpretable understanding of treatment outcomes on vaginal microbiota. Antibiotic treatment led to sparser networks in no-relapse group, reflecting potential disruption of dysbiotic co-occurrence patterns and movement toward a healthy equilibrium (Figure 4 and 5). BV was characterized by densely connected microbial networks, dominated by associations between pathogenic bacteria. Persistence of *Sneathia* spp. in post-treatment low-lactobacilli dominant environments often co-occurring with other pro-inflammatory bacteria was notable in relapse state. *Sneathia sanguinegens* particularly remained a central node in both baseline and relapse networks, suggesting its role as a keystone taxon that may resist displacement and drive recurrence. Its consistent negative association with *Lactobacillus* and positive correlation with *Gardnerella* and *Dialister* further implied antagonistic relationships that could help maintain a dysbiotic state hindering re-colonization by beneficial microbes. Several studies that report co-occurrence of *S. sanguinegens* and *G. vaginalis* have in fact discussed lower treatment success rates.^16,27^ These findings consolidate *S. sanguinegens* as a key target for future studies and potentially precision interventions aimed at preventing BV recurrence.

Emergence of *Corynebacterium amycolatum* as a prominent node in the microbial community network in no-relapse was noteworthy. While traditionally underexplored in the context of vaginal health, emerging studies suggest *C. amycolatum* has the potential to produce antimicrobial substances.(30) Its presence in no-relapse, may hold potential significance in aiding re-establishment of microbial homeostasis, preventing pathogen growth and favouring re-colonization of *Lactobacillus* spp. Altogether, the observed patterns of shifts and persistence in network composition, density and membership, especially identification of marker taxa like *S. sanguinegens, Dialister* and *C. amycolatum*, provided interpretable indicators of treatment outcomes in BV. It would be prudent to acknowledge the challenges and limitations of the study. Given the social stigma associated with sexual health in low-income countries, the self-reporting of the disorder and challenges of follow-up, inaccuracy in recall of extended medical and personal hygiene history may have occurred, potentially influencing the reliability of the collected data. Further, given the low-no-relapse or recovery rate, the small sample size of the no-relapse group underpowers statistical comparisons and generalizability, warranting careful interpretations. While trends in microbial shifts and network transitions are compelling, longitudinal studies with larger patient cohorts are thus necessary to validate the identified microbial indicators and their predictive value for treatment outcomes. Interpretation of taxa-specific roles and their contributions to community resilience or recovery through functional assays or metagenomic approaches would further strengthen these findings.

## Acknowledgements

The authors also acknowledge Indian Council of Medical Research, Government of India for research fellowship award (5/3/8/35/ITR-F/2019-ITR) to Dr. Apoorva Challa.

## References

1. Atashili J, Poole C, Ndumbe PM, Adimora AA, Smith JS. Bacterial vaginosis and HIV acquisition: a meta-analysis of published studies. AIDS. 2008;22(12):1493–1501. doi:10.1097/QAD.0b013e3283021a37

2. Ness RB, Hillier SL, Kip KE, et al. Bacterial vaginosis and risk of pelvic inflammatory disease. Obstet Gynecol. 2004;104(4):761–769. doi:10.1097/01.AOG.0000139512.37582.17

3. Leitich H, Kiss H. Asymptomatic bacterial vaginosis and intermediate flora as risk factors for adverse pregnancy outcome. Best Pract Res Clin Obstet Gynaecol. 2007;21(3):375–390. doi:10.1016/j.bpobgyn.2006.12.005

4. Workowski KA, Bachmann LH, Chan PA, et al. Sexually transmitted infections treatment guidelines, 2021. MMWR Recomm Rep. 2021;70(4):1–187. doi:10.15585/mmwr.rr7004a1

5. Bradshaw CS, Morton AN, Hocking J, et al. High recurrence rates of bacterial vaginosis over the course of 12 months after oral metronidazole therapy and factors associated with recurrence. J Infect Dis. 2006;193(11):1478–1486. doi:10.1086/503780

6. Muzny CA, Kardas P. A narrative review of current challenges in the diagnosis and management of bacterial vaginosis. Sex Transm Dis. 2020;47(7):441–446. doi:10.1097/OLQ.0000000000001178

7. Ravel J, Gajer P, Abdo Z, et al. Vaginal microbiome of reproductive-age women. Proc Natl Acad Sci U S A. 2011;108(Suppl 1):4680–4687. doi:10.1073/pnas.1002611107

8. Oluwatosin Goje, Elizabeth O. Shay, Metabel Markwei, Roshan Padmanabhan, Charis Eng. The effect of oral Metronidazole on the vaginal microbiome of patients with recurrent bacterial vaginosis: A pilot investigational study. Human Microbiome Journal. 20; 2021. 10.1016/j.humic.2021.100081.

9. Swidsinski A, Mendling W, Loening-Baucke V, et al. Adherent biofilms in bacterial vaginosis. Obstet Gynecol. 2005;106:1013–1023. doi:10.1097/01.AOG.0000183594.45524.d2

10. Challa A, Maras JS, Nagpal S, et al. Multi-omics analysis identifies potential microbial and metabolite diagnostic biomarkers of bacterial vaginosis. J Eur Acad Dermatol Venereol. 2024 Jun;38(6):1152–1165. doi: 10.1111/jdv.19805. Epub 2024 Jan 29. PMID: 38284174.

11. Nagpal S, Singh R, Yadav D, Mande SS. MetagenoNets: comprehensive inference and meta-insights for microbial correlation networks. Nucleic Acids Res. 2020 Jul 2;48(W1):W572–W579. doi: 10.1093/nar/gkaa254. PMID: 32338757; PMCID: PMC7319469.

12. Nagpal S, Kuntal BK, Mande SS. NetSets.js: a JavaScript framework for compositional assessment and comparison of biological networks through Venn-integrated network diagrams. Bioinformatics. 2021 May 1;37(4):580–582. doi: 10.1093/bioinformatics/btaa723. PMID: 32805035.

13. https://graphonline.top/

14. Shannon P, Markiel A, Ozier O, et al. Cytoscape: a software environment for integrated models of biomolecular interaction networks. Genome Res. 2003 Nov;13(11):2498–504. doi: 10.1101/gr.1239303. PMID: 14597658; PMCID: PMC403769.

15. Petrina MAB, Cosentino LA, Rabe LK, Hillier SL. Susceptibility of bacterial vaginosis (BV)-associated bacteria to secnidazole compared to metronidazole, tinidazole and clindamycin. Anaerobe. 2017 Oct;47:115–119. doi: 10.1016/j.anaerobe.2017.05.005. Epub 2017 May 15. PMID: 28522362; PMCID: PMC5623164.

16. Gustin AT, Thurman AR, Chandra N, et al. Recurrent bacterial vaginosis following metronidazole treatment is associated with microbiota richness at diagnosis. Am J Obstet Gynecol. 2022;226(2):225.e1–225.e15. doi:10.1016/j.ajog.2021.09.018

17. Vodstrcil LA, Muzny CA, Plummer EL, Sobel JD, Bradshaw CS. Bacterial vaginosis: drivers of recurrence and challenges and opportunities in partner treatment. BMC Med. 2021;19(1):194. Published 2021 Sep 2. doi:10.1186/s12916-021-02077-3

18. Kim MJ, Lee S, Kwon MY, Kim M. Clinical Significance of Composition and Functional Diversity of the Vaginal Microbiome in Recurrent Vaginitis. Front Microbiol. 2022;13:851670. Published 2022 Feb 18. doi:10.3389/fmicb.2022.851670

19. Ceccarani, C., Foschi, C., Parolin, C. et al. Diversity of vaginal microbiome and metabolome during genital infections. Sci Rep 9, 14095 (2019). 10.1038/s41598-019-50410-x

20. Pendharkar S, Skafte-Holm A, Simsek G, Haahr T. Lactobacilli and Their Probiotic Effects in the Vagina of Reproductive Age Women. Microorganisms. 2023 Mar 1;11(3):636. doi: 10.3390/microorganisms11030636. PMID: 36985210; PMCID: PMC10056154.

21. Fredricks DN, Plantinga A, Srinivasan S, et al. Vaginal and Extra-Vaginal Bacterial Colonization and Risk for Incident Bacterial Vaginosis in a Population of Women Who Have Sex With Men. J Infect Dis. 2022 Apr 1;225(7):1261–1265. doi: 10.1093/infdis/jiaa233. PMID: 32379324; PMCID: PMC8974833.

22. Theis KR, Florova V, Romero R, Borisov AB, Winters AD, Galaz J, Gomez-Lopez N. Sneathia: an emerging pathogen in female reproductive disease and adverse perinatal outcomes. Crit Rev Microbiol. 2021 Aug;47(4):517–542. doi: 10.1080/1040841X.2021.1905606. Epub 2021 Apr 6. PMID: 33823747; PMCID: PMC8672320.

23. Mendling W, Palmeira-de-Oliveira A, Biber S, Prasauskas V. An update on the role of Atopobium vaginae in bacterial vaginosis: what to consider when choosing a treatment? A mini review. Arch Gynecol Obstet. 2019 Jul;300(1):1–6. doi: 10.1007/s00404-019-05142-8. Epub 2019 Apr 5. PMID: 30953190; PMCID: PMC6560015.

24. Morio F, Jean-Pierre H, Dubreuil L, et al. Antimicrobial susceptibilities and clinical sources of Dialister species. Antimicrob Agents Chemother. 2007;51(12):4498–4501. doi:10.1128/AAC.00538-07

25. Marrazzo JM, Thomas KK, Fiedler TL, Ringwood K, Fredricks DN. Relationship of specific vaginal bacteria and bacterial vaginosis treatment failure in women who have sex with women. Ann Intern Med. 2008 Jul 1;149(1):20–8. doi: 10.7326/0003-4819-149-1-200807010-00006. PMID: 18591634; PMCID: PMC2630802.

26. Mollin A, Katta M, Sobel JD, Akins RA. Association of key species of vaginal bacteria of recurrent bacterial vaginosis patients before and after oral metronidazole therapy with short- and long-term clinical outcomes. PLoS One. 2022 Jul 28;17(7):e0272012. doi: 10.1371/journal.pone.0272012. PMID: 35901180; PMCID: PMC9333308.

27. Srinivasan S, Hoffman NG, Morgan MT, et al. Bacterial communities in women with bacterial vaginosis: high resolution phylogenetic analyses reveal relationships of microbiota to clinical criteria. PLoS One. 2012;7(6):e37818. doi:10.1371/journal.pone.0037818

28. Amorim AT, Lino VS, Marques LM, et al. Mycoplasma hominis Causes DNA Damage and Cell Death in Primary Human Keratinocytes. Microorganisms. 2022;10(10):1962. Published 2022 Oct 1. doi:10.3390/microorganisms10101962

29. Chew J, Zilm PS, Fuss JM, Gully NJ. A proteomic investigation of Fusobacterium nucleatum alkaline-induced biofilms. BMC Microbiol. 2012;12:189. Published 2012 Sep 3. doi:10.1186/1471-2180-12-189

30. Gladysheva IV, Cherkasov SV, Khlopko YA, Plotnikov AO. Genome Characterization and Probiotic Potential of Corynebacterium amycolatum Human Vaginal Isolates. Microorganisms. 2022;10(2):249. Published 2022 Jan 23. doi:10.3390/microorganisms10020249

