## Supplementary figures and images for "Tracking vaginal microbiome transitions in bacterial vaginosis for cues of antibiotic resilience"

### Figure S1

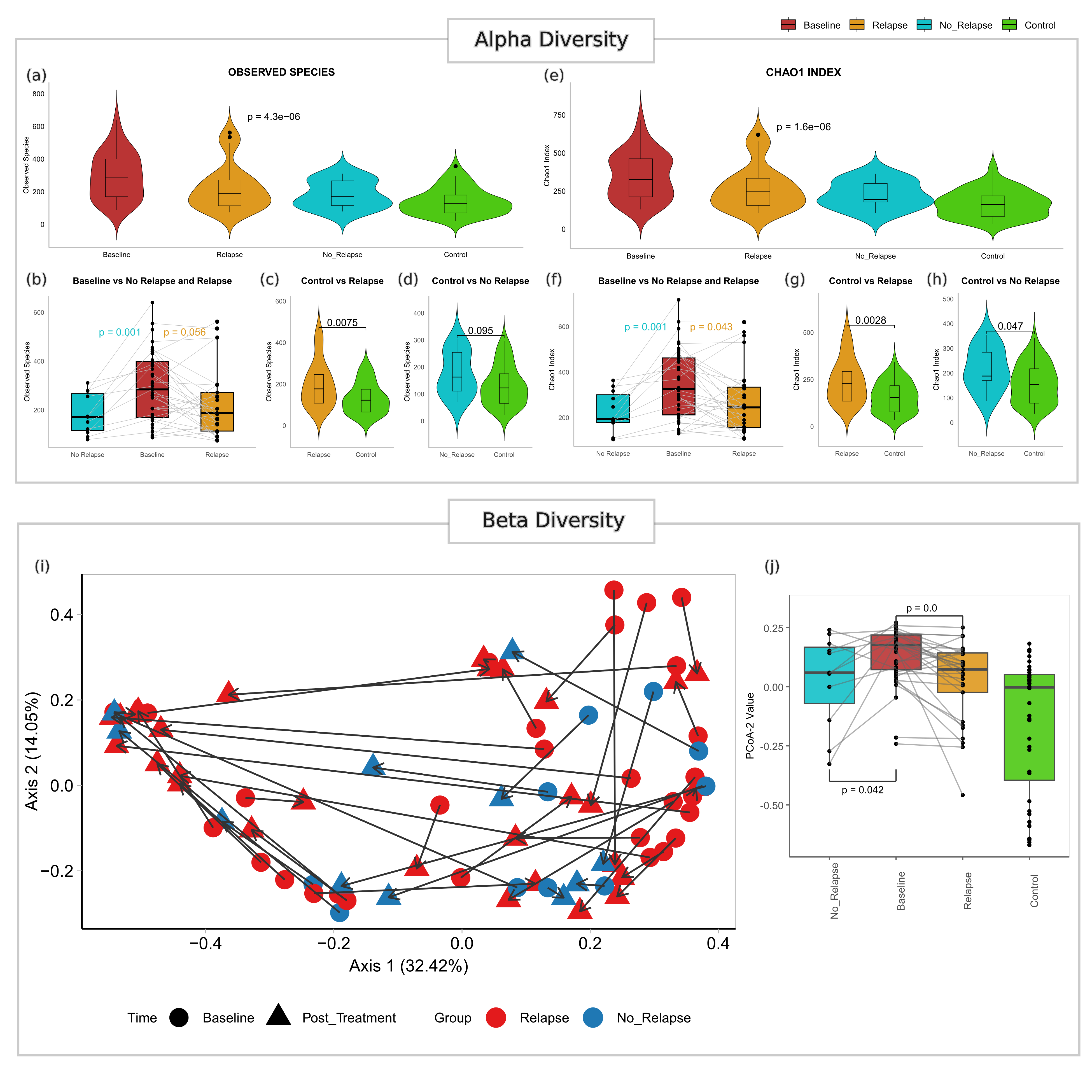

### Figure S2

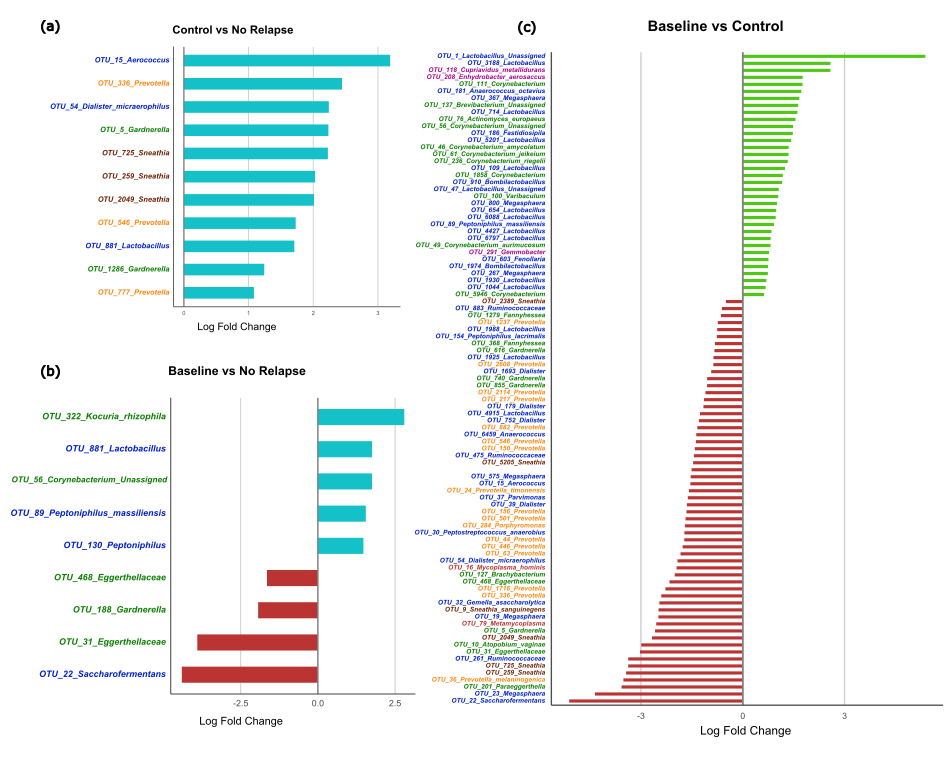

### Figure S3

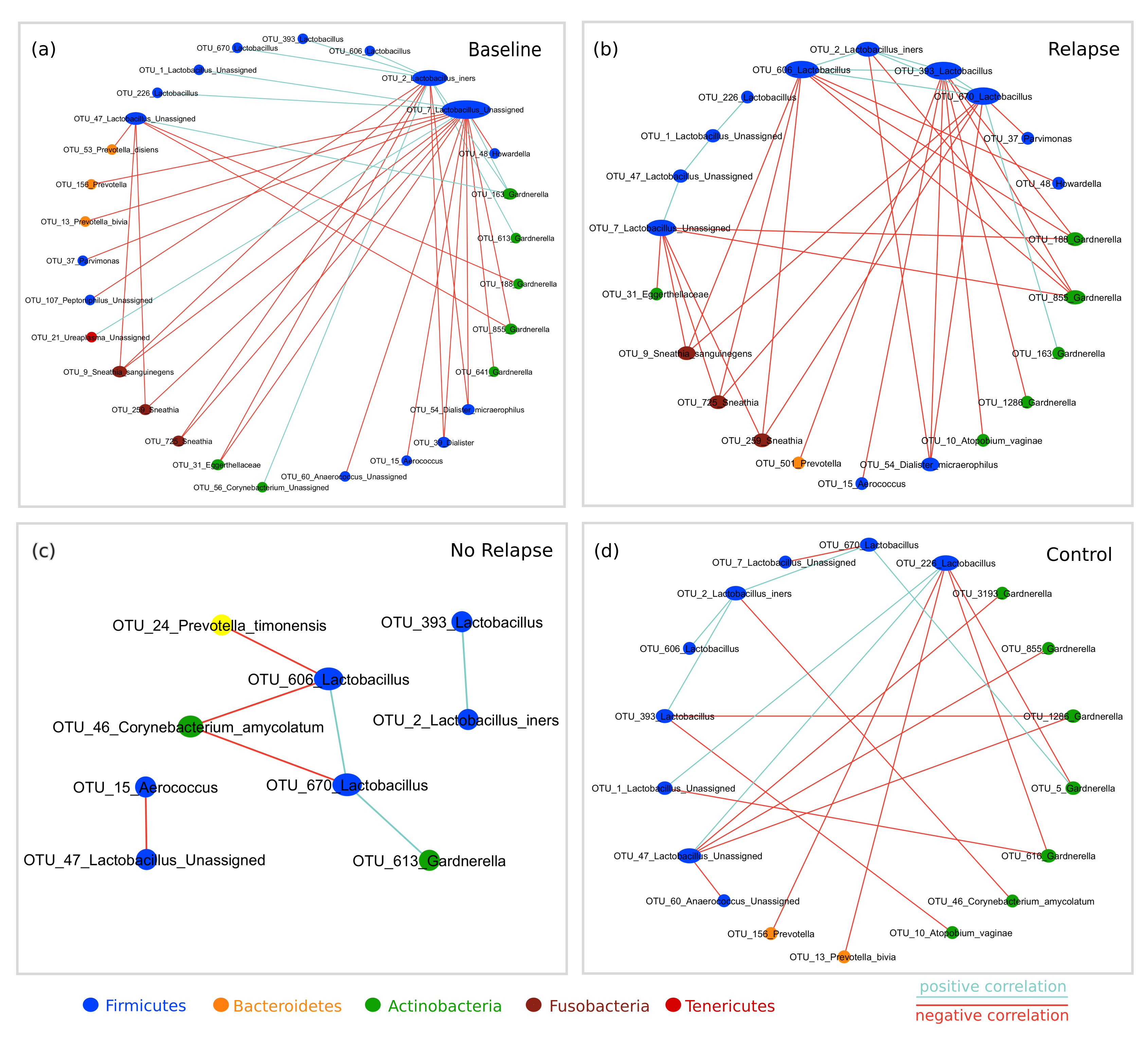

### Figure S4

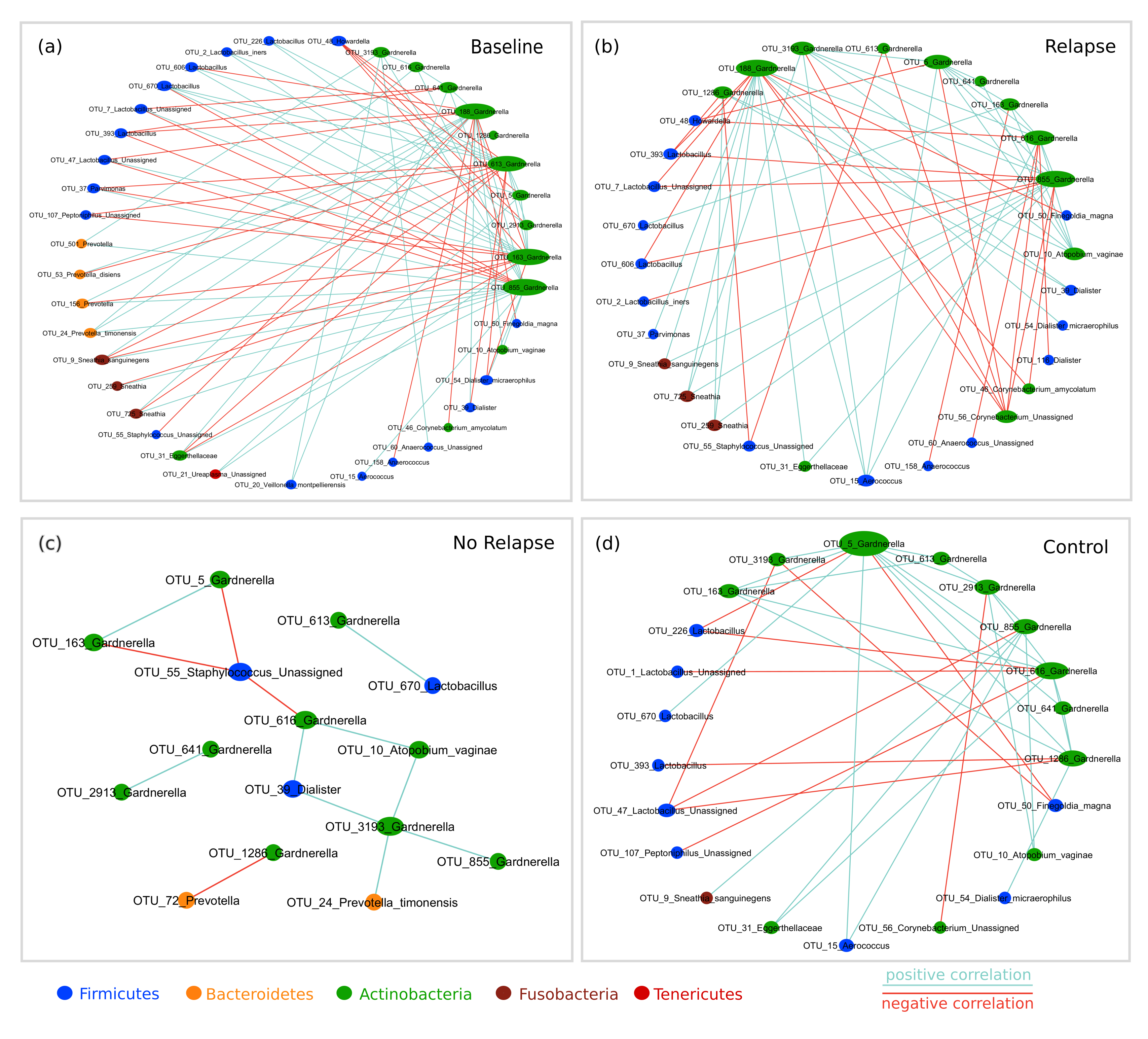

### Figure S5

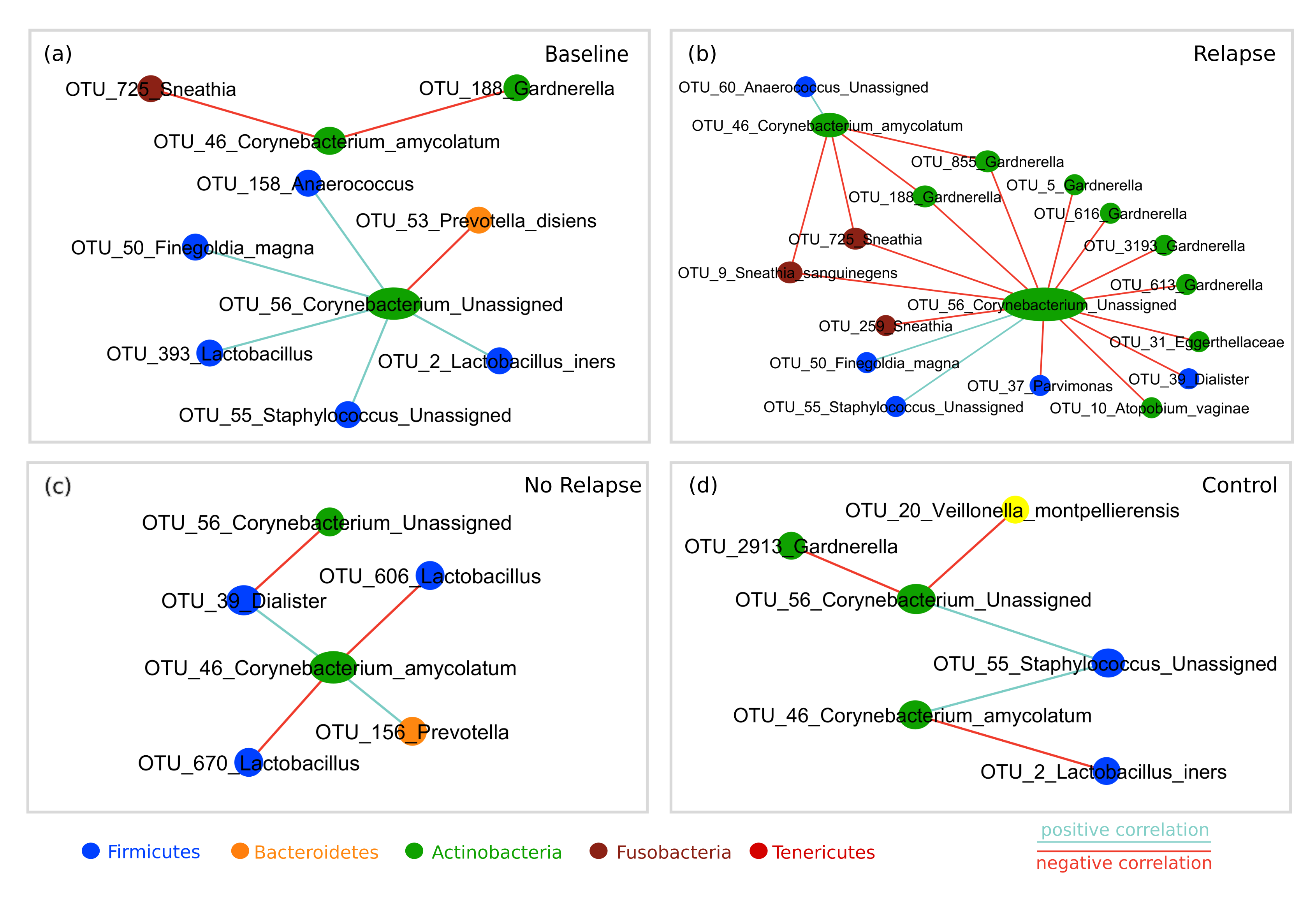
